# Liver size is predetermined in the neonate by adding lobules at the periphery

**DOI:** 10.1101/2023.10.12.562109

**Authors:** D. Berfin Azizoglu, Karina Perez, Sherry Li Zheng, Shahadat Rahman, Ellen Youngsoo Rim, Teni Anbarchian, Matt Fish, Kyle M. Loh, Kristy Red-Horse, Roel Nusse

## Abstract

Organs vary in size between and within species to match organismal needs^1,2^. Decades-old theoretical work has proposed that scaling of organs and body parts relative to the body relies on the features of energy-transport systems, the vascular system in mammals^3^. Yet, experimental studies on whether or how vascularization helps determine organ size have lagged behind. The mammalian liver is a remarkable example, as liver size scales proportionally with high precision between individuals^4^. Here, we use quantitative clonal mapping, volumetric imaging, and genetic perturbations combined with novel molecular and genetic tools to identify the temporal and spatial constraints that establish mouse liver size. We find that adult liver size is predetermined during a neonatal period when new functional units, termed lobules, are added to the organ. New lobules are vascularized by prominent sprouting angiogenesis of the hepatic vein, restricted to the periphery of the organ. When Wnt signals are ablated in the single cell-layered mesothelium at the periphery, lobule growth fails, and the organ adopts a compromised size set point. Remarkably, within a week after birth and well before hepatocyte division stops, vein sprouting rapidly declines and lobule addition concludes, setting a limit on the final liver size. These findings posit that vascularization in the neonate constrains and helps determine adult liver size. Together, these results propose a novel, vasculature-centric experimental framework for studying organ size control and scaling in mammals.

## Main

Organ size scales with animal size in mammals that themselves vary up to a hundred million times in weight^2^. How mammalian organs reach their proper size across such a wide range of scales despite similar developmental trajectories is a fundamental question in biology with direct implications for regenerative medicine^5,6^. Our understanding of organ size control has largely relied on studies of parenchymal cell proliferation and the relevant signaling networks^2^. However, parenchymal cells alone cannot support an organ; multiple cell types must integrate into anatomical units for an organ to form and function^7^. Indeed, one central regulator of parenchymal cell proliferation, Hippo signaling, fails to instruct architecturally normal organ growth on its own^8–11^. Theoretical modeling from the 1990s instead suggests that energy-distributing vascular systems are a key component of scaling body parts relative to the body^3^. These observations highlight the need to study organ size in the context of multicellular integration and, in particular, vascularization. Here, we focus on the mouse liver to interrogate the multicellular-scale mechanisms that determine mammalian organ size. Liver weight is set at a stereotyped ratio to body weight, offering a tractable model system^1,2,4^. The adult liver consists of repeated functional units termed lobules^12^. Thus, proper liver size must be achieved by fine-tuning the number and size of lobules in a given individual. We investigate the mechanisms that establish the liver size by studying how new liver lobules form and vascularize, and, importantly, cease to form.

### Novel liver lobule marker reveals generation of lobules after birth

To study how lobule organization relates to liver size in mice, we sought to identify a molecular marker of mouse liver lobules. Demarcating liver lobules in rodents and humans has proven challenging, as the extracellular matrix components at lobule boundaries are scarce in these species unlike in other mammals^12,13^. We found that a component of the coagulation pathway, Protein C Receptor (Procr), is highly enriched in the portal veins and venules (referred to as portal vessels) and thus outlines liver lobules (Fig. 1a-a’’, Extended Data Fig. 1a). To define when lobule boundaries are assembled, we assessed Procr expression through the postnatal period (Fig. 1b-c’). Lobule boundaries were discernable as early as one week of age (Fig. 1c-c’, Extended Data Fig. 1b-d’). These results establish Procr as a novel marker of lobule boundaries and suggest that the mouse liver is organized into lobules neonatally.

**Fig. 1.**
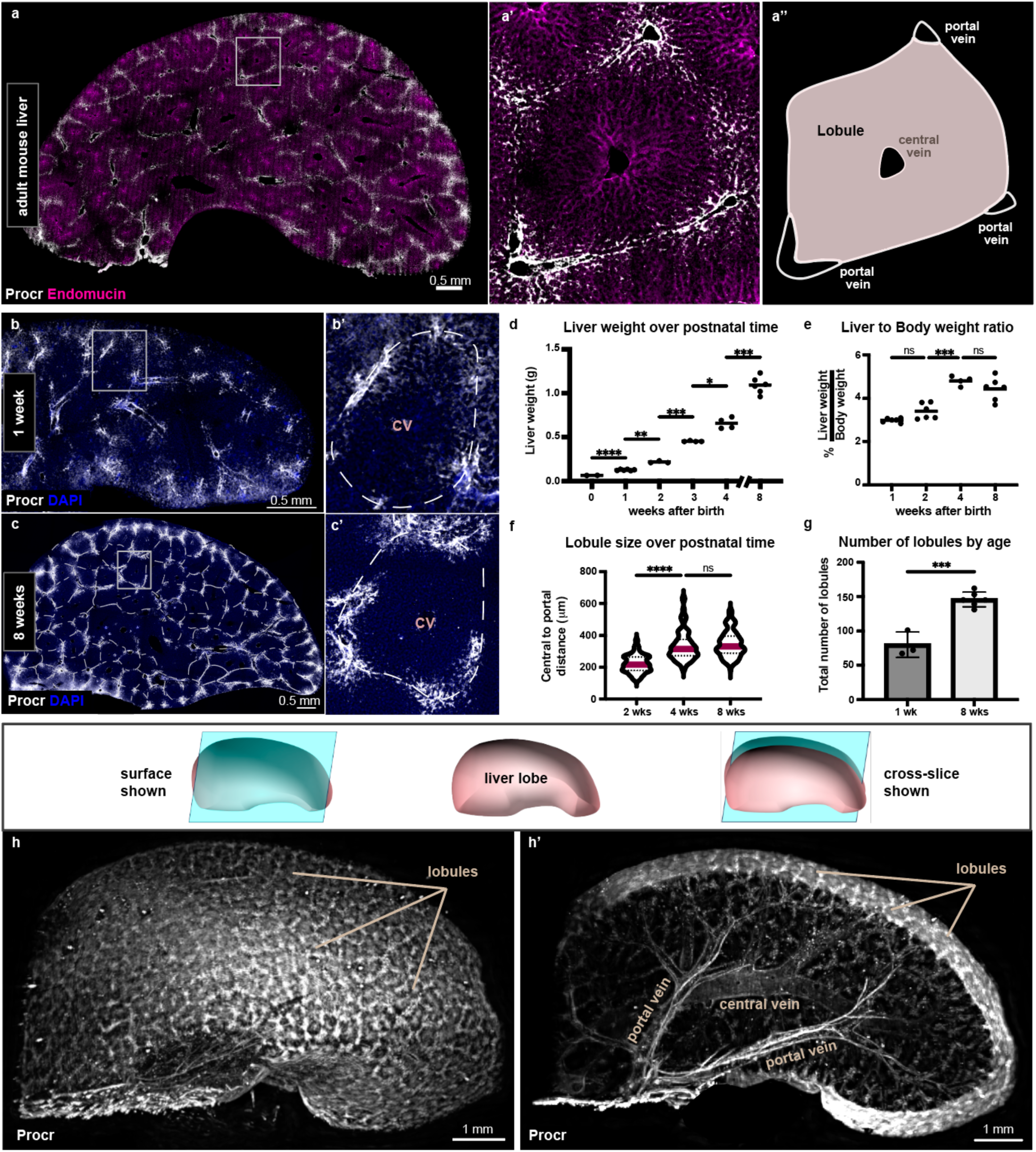
Novel liver lobule marker reveals generation of lobules after birth. **a-a’**, Thick section of the adult mouse liver immunostained for the lobule boundary marker Procr and the venous endothelial marker Endomucin. **a’’,** Schematic representation of the lobule. **b-c’,** Thick section of the liver immunostained for the lobule boundary marker Procr from 1 week- and 8 week-old mice. Dotted lines delineate lobules. **d,** Increase in absolute liver weight over postnatal time for n=2 newborns, n=6 (1 week), n=3 (2 weeks), n=4 (3 weeks, male), n=4 (4 weeks, male), and n=6 (8 weeks, male) mice. Brown-Forsythe Anova test and Dunnett’s multiple comparisons test performed. For clarity, differences between consecutive ages are shown. Adjusted p-values from left to right: *< 0.0001, 0.0072, 0.0004, 0.0451, 0.0003*. **e,** Liver to body weight ratio as percentage for n=6 (1 week), n=6 (2 weeks), n=4 (4 weeks, male), and n=6 (8 weeks, male) mice. Brown-Forsythe Anova test and Dunnett’s multiple comparisons test performed. For clarity, differences between consecutive ages are shown. Adjusted p-values from left to right: *0.2165, 0.0005, 0.7221*. **f,** Violin plot shows the distance from the central vein to portal vessels at lobule boundaries in livers of 2 week-, 4 week-, and 8-week-old mice. Red bars represent the mean. A total of n=210 (2 weeks), n=147 (4 weeks), and n=162 (8 weeks) lobules were scored in 3 mice per group. Brown-Forsythe Anova test and Dunnett’s multiple comparisons test, adjusted p-values from left to right: *< 0.0001, > 0.9999*. **g,** The total number of lobules in one whole liver lobe of 1 week-old (n=3) and 8-week-old (n=6) mice. Mean values ± s.d. shown. Unpaired t test, *p-value = 0.0002*. **h-h’,** Volumetric images of whole liver lobes iDisco-cleared and immunostained for the lobule boundary marker Procr are shown in surface view in **h** and slice view in **h’**. ns, not significant. DAPI in blue depicts nuclei.

Next, we sought to define when the stereotyped ratio of liver to body weight, the liver size set point, is achieved. While the mouse liver grows in absolute weight from birth until eight weeks of age (Fig. 1d), we found that it reaches its stereotyped size set point well before that, at four weeks of age (Fig. 1e). This prompted us to examine how lobule size and numbers change over the same period. Lobules became larger over postnatal time (Fig. 1f). Strikingly, lobule size was finalized by four weeks of age (Fig. 1f), long prior to the absolute liver weight, and concurrently with the liver to body weight ratio (Fig. 1e-f). These results couple lobule size to the liver size set point.

To determine potential changes in lobule numbers, we assessed the total number of lobules in whole liver lobes. To this end, we optimized a method that uses iDisco clearing^14^ combined with Procr immunostaining and light-sheet imaging (Fig. 1g-h’, Extended Data Video 1, 2). Lobules could be seen covering the entire surface of the imaged livers (Fig. 1h). Procr labeling enabled us to simultaneously visualize the portal and central vein branches feeding into the lobules deep through the organ (Fig. 1h’). Surprisingly, quantification of mature lobules revealed a twofold increase in lobule number over the postnatal period (Fig. 1g), indicating that close to half of adult liver lobules are added to the organ postnatally.

These findings suggest that the adult liver size set point is established postnatally through lobule enlargement and, unexpectedly, the formation of new lobules.

### Cells divide and expand dramatically in lobules at the organ periphery

Lobule size is highly stereotyped within species and therefore cannot explain intra-species variation in liver size^12^. By contrast, lobule numbers may vary among individuals and contribute to the establishment of the liver size set point. We therefore sought to understand what factors constrain lobule numbers in a given liver. To address this, we tracked the formation of new lobules in the postnatal liver.

We reasoned that adding a new lobule would require substantial cell proliferation. To survey the extent of proliferation, we measured the frequency of mitoses in full liver sections by phospho-histone H3 (pHH3) labeling. This analysis identified a peak of cell division at postnatal day (P) 8, immediately after the first postnatal week (Fig. 2a). Spatial distribution of the mitotic cells was surprisingly uneven, with a strong enrichment towards the edge of the liver (Fig. 2b-c). This indicated that more cell division occurs at the edge of the liver early in the postnatal period.

**Fig. 2.**
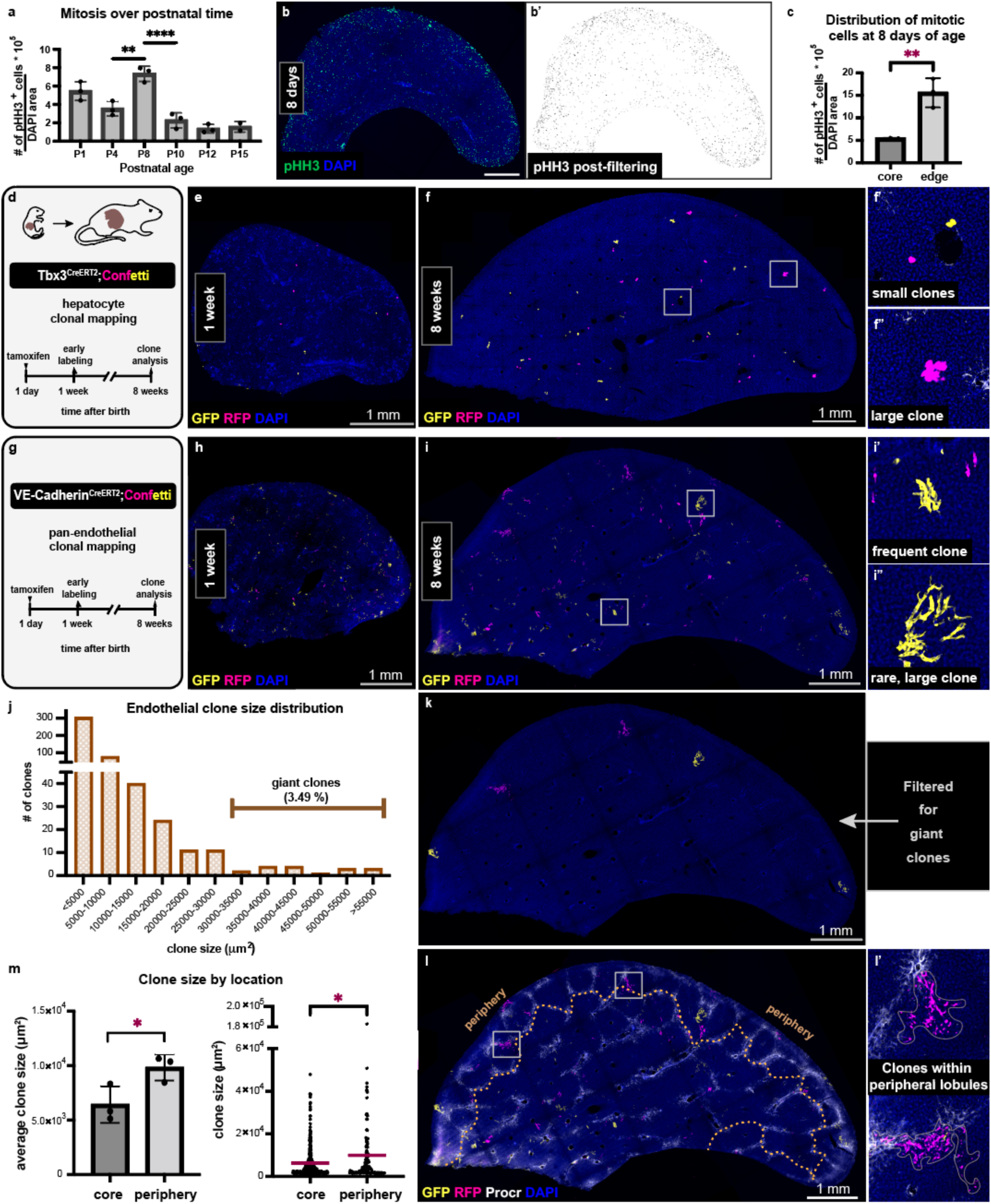
Cells expand dramatically in a subset of liver lobules at the organ periphery. **a**, Quantification of phospho-histone H3 (pHH3)+ cells at postnatal ages (P). Mean values ± s.d. n=3 mice per group except n=2 mice at P15. Ordinary one-way Anova analysis and Tukey’s multiple comparisons test performed. For clarity, significant differences between consecutive ages are shown. Adjusted p-values from left to right, including the non-significant comparisons between consecutive ages: *0.0962, 0.001, < 0.0001, 0.7127, 0.9996*. **b,** Thick liver section at 8 days immunostained for pHH3. **b’,** Post-processing of image in **b** with intensity thresholding and particle analysis. **c,** Quantification of pHH3+ cells at the edge and the core at 8 days from n=3 mice. Mean values ± s.d. Unpaired t test, *p-value = 0.0054*. **d,** Schematic summary of the experimental design to map hepatocyte expansion. **e-f’’,** Tbx3^CreERT2^;Confetti thick liver section immunostained for GFP and RFP to visualize the early labeling at 1 week and the clones at 8 weeks. **g,** Schematic summary of the experimental design to map endothelial cell expansion. **h-i’’,** VE-Cadherin^CreERT2^;Confetti thick liver section immunostained for GFP and RFP to visualize the early labeling at 1 week and the clones at 8 weeks. **j,** Endothelial clone size distribution from VE-Cadherin^CreERT2^;Confetti livers at 8 weeks, n=457 clones from 3 mice. **k,** Clones in **i** filtered by size (>3×10^4^ µm^2^) to visualize only the giant clones. **l-l’,** Merge of image in **i** with Procr immunostaining with the periphery delineated (dotted line). **m,** Comparison of endothelial clone size between the core and the periphery in 8-week-old mice. Mean values ± s.d. per mouse (left) and individual clone values (right). n=128 clones in the periphery and n=329 clones in the core from 3 mice. Left: Ratio paired t test, *p-value = 0.0417*. Right: Welch’s t test, *adjusted p-value = 0.036*. DAPI in blue depicts nuclei. tam, tamoxifen.

We asked whether the uneven distribution of cell division translates to biased growth over the postnatal period. We monitored the long-term expansion of cells in the liver, mainly hepatocytes and cells of the hepatic vasculature. To assess hepatocyte expansion, we performed lineage tracing using an inducible Tbx3^Confetti^ mouse model where Tbx3^CreERT2^ drives the Confetti reporter upon Tamoxifen administration (Fig. 2d). Tbx3 is neonatally expressed in many hepatocytes except those in the periportal zone^15^ (Extended Data Fig. 1a). We sparsely labeled the cells at one day of age and confirmed that, by one week of age, hepatocytes were labeled across the liver (Fig. 2e). By adulthood, many hepatocytes had formed multicellular clones (Fig. 2f). The clones were distributed across the liver, suggesting that growth occurred across the organ. However, we noted that particularly large clones had emerged towards the edge of the liver (Fig. 2f-f’’, Extended Data Fig. 2a).

We next assessed the expansion of another major liver cell type, vascular endothelial cells, as vascularization is indispensable for the growth of the organ^16–19^. We targeted endothelial cells across vessel subtypes, i.e., veins, arteries, and capillaries, by combining the tamoxifen-inducible pan-endothelial VE-Cadherin^CreERT2^ with the Confetti reporter system (VeCad^Confetti^). We sparsely labeled the cells at one day of age and performed clonal analysis at adulthood (Fig. 2g). Labeled endothelial cells were detectable throughout the liver one week following tamoxifen induction (Fig. 2h), indicating that labeling occurs efficiently across the organ. By adulthood, most single cells expanded into multicellular clones (Fig. 2i-i’’). Similar to hepatocytes, endothelial cells expanded across the liver, but larger endothelial clones appeared towards the edge (Fig. 2i-i’’). Intriguingly, rare clones over 30,000 µm^2^ in area emerged which we refer to as giant clones (Fig. 2j). These giant clones strictly localized towards the edge of the liver (Fig. 2k).

The co-occurrence of larger endothelial and hepatocyte clones suggested accelerated growth at the edge of the organ. To test if this growth underpins new lobule formation, we mapped the endothelial clones against Procr-marked lobule boundaries. This uncovered a striking relationship: the giant clones were 4 times more likely to reside within the outermost lobules at the edge of the liver, referred to as the periphery (Fig. 2l-l’, Extended Data Fig. 2a). Endothelial and hepatocyte clones in the periphery defined by lobule boundaries were significantly larger on average compared to the clones in the core (Fig. 2m, Extended Data Fig. 2b). We confirmed that this is not due to differences in labeling efficiency (Extended Data Fig. 2c). These results demonstrate that peripheral lobules grow extensively and are likely the youngest lobules.

Considering the increase in lobule number over the postnatal period, these findings suggest that the peripheral liver lobules are newly formed after birth.

### Mesothelial Wnts promote lobule growth and fine-tune adult liver size

We hypothesized that the liver periphery must contain a source of growth signals to promote the development of new lobules. A single epithelial layer, the mesothelium, encapsulates the liver at its periphery and is a known source of mitogens in the embryo^20,21^. Among these mitogens, Wnts are highly expressed in the mesothelium^21^ and are well-known regulators of liver growth^22^. Indeed, the postnatal hepatic mesothelium expressed high levels of Wntless which is required for Wnt secretion (Fig. 3a-a’), motivating us to test the role of mesothelial Wnt signals. We generated mice with mesothelial-specific, inducible Wntless loss of function, WT1^CreERT2^; Wntless^f/f^ (Wntless^meso^) (Fig. 3b, Extended Data Fig. 3a-b’). Depletion of mesothelial Wntless in neonates was sufficient to dampen the peripheral enrichment of mitosis (Fig. 3c-d, Extended Data Fig. 3c). While mitotic activity in control livers was concentrated to the periphery, mesothelial Wntless deletion led to a more even distribution of mitotic cells across the liver. Remarkably, by adulthood, this resulted in a significant shift in liver size set point. In Wntless^meso^ mice, the liver-to-body weight ratio was lower compared to controls (Fig. 3e, Extended Data Fig. 3d-e). This was a consequence of impaired lobule growth, as Wntless^meso^ lobule size was significantly reduced (Fig. 3f). Wntless^meso^ mice showed no gross abnormalities in liver architecture, lobule organization (Fig. 3g-g’’), or the number of lobules (Extended Data Fig. 3f).

**Fig. 3.**
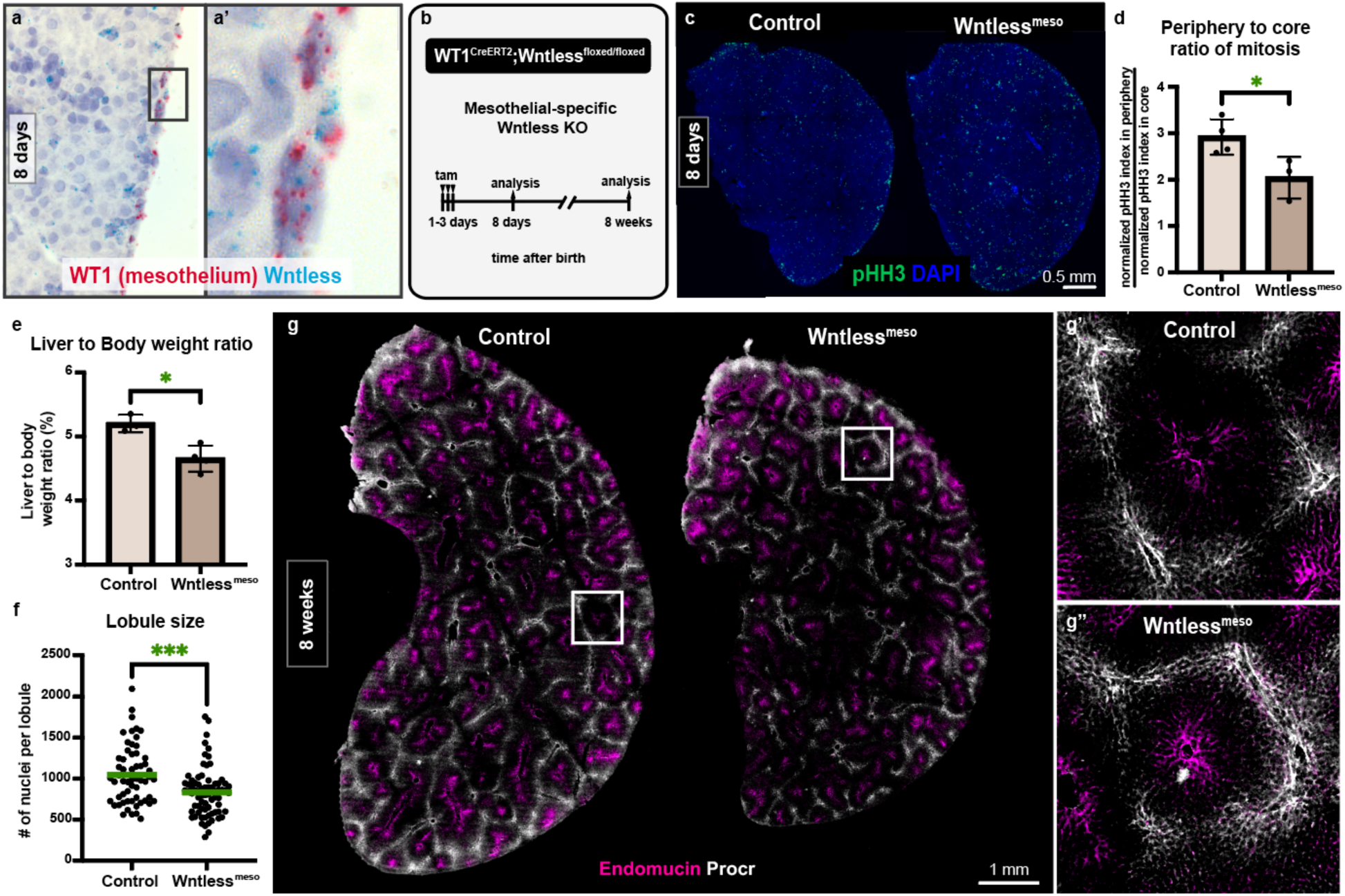
Liver size set point relies of peripheral tissue expansion promoted by mesothelial Wnt signaling. **a-a’**, In situ hybridization on a representative wildtype liver section for *Wntless* and the mesothelial-specific *Wilms Tumor 1 (WT1)* at 8 days. **b,** Schematic summary of the experimental design to determine the role of mesothelial Wnt signaling in postnatal growth and size control of the liver. tam, tamoxifen. **c,** Thick left lobe liver sections of control and mesothelial-specific Wntless knock-out (KO), or Wntless^meso^, mice at 8 days old immunostained for phospho-histone H3. **d,** Quantification of mitosis enrichment in the liver periphery for 8 days old control (n=4) and mesothelial-specific Wntless knock-out (KO) (n=3) mice measured as the ratio of mitotic index in the periphery to that in the core. Mean values ± s.d. Unpaired t test, *p-value = 0.0376.* **e,** Liver to body weight ratio of 8-week-old female control (n=3) and mesothelial-specific Wntless knock-out (KO) (n=4) mice. Mean values ± s.d. Unpaired t test, *p-value = 0.0103*. **f,** Lobule size measured as the number of nuclei per lobule in 8-week-old female control (n=58 lobules in 2 mice) and mesothelial-specific Wntless KO (n=73 lobules in 2 mice). Green bars represent the mean values. Unpaired t test, *p-value = 0.0003*. **g-g’’,** Thick liver sections of female control and mesothelial-specific Wntless KO mice immunostained for the venous marker Endomucin and the lobule boundary marker Procr. **g’** and **g’’** show single lobules from each genotype. DAPI in blue depicts nuclei.

Together, these results demonstrate that mesothelial Wnts are necessary for liver lobule growth and identify the mesothelium as an unexpected organizer of postnatal liver growth. Importantly, these findings suggest that biased growth at the periphery is required to establish the proper adult liver size set point.

### Peripheral vascularization of venous origin demarcates newly formed lobules

We next asked how new lobules at the periphery initiate and become vascularized. Vascularization is likely to reveal key aspects of new lobule formation, as blood vessels provide reliable landmarks in the lobule^12,13^. To study lobule vascularization, we first needed to gain genetic access to hepatic vessel subtypes. Thus, we surveyed a series of tamoxifen-inducible vascular endothelial Cre mouse lines (Extended Data Fig. 4). Among these, two lines were successful at targeting the large vessels of the lobule: Apelin^CreERT2^ that labeled the central veins (Extended Data Fig. 4b), and Efnb2^CreERT2^ that labeled the portal vessels (Extended Data Fig. 4c). By crossing these lines into the Confetti or the mTmG reporter, we induced labeling soon after birth (Fig. 4a, d) and assessed the contribution of each vessel type to the vascularization of new lobules at the periphery. Endothelial cells of portal vessels formed clones as seen with Efnb2^Confetti^ tracing (Extended Data Fig. 5a-b); these clones were similarly sized and not spatially biased (Extended Data Fig. 5a’-b). By contrast, central vein endothelial cells marked by Apelin^Confetti^ formed clones of highly varying sizes (Fig. 4b, b’’), evidently larger in the periphery than in the core (Fig. 4b-c). These data demonstrated that central vein cells expand preferentially at the periphery, raising the possibility that central veins serve as origins of vasculature in new lobules.

**Fig. 4.**
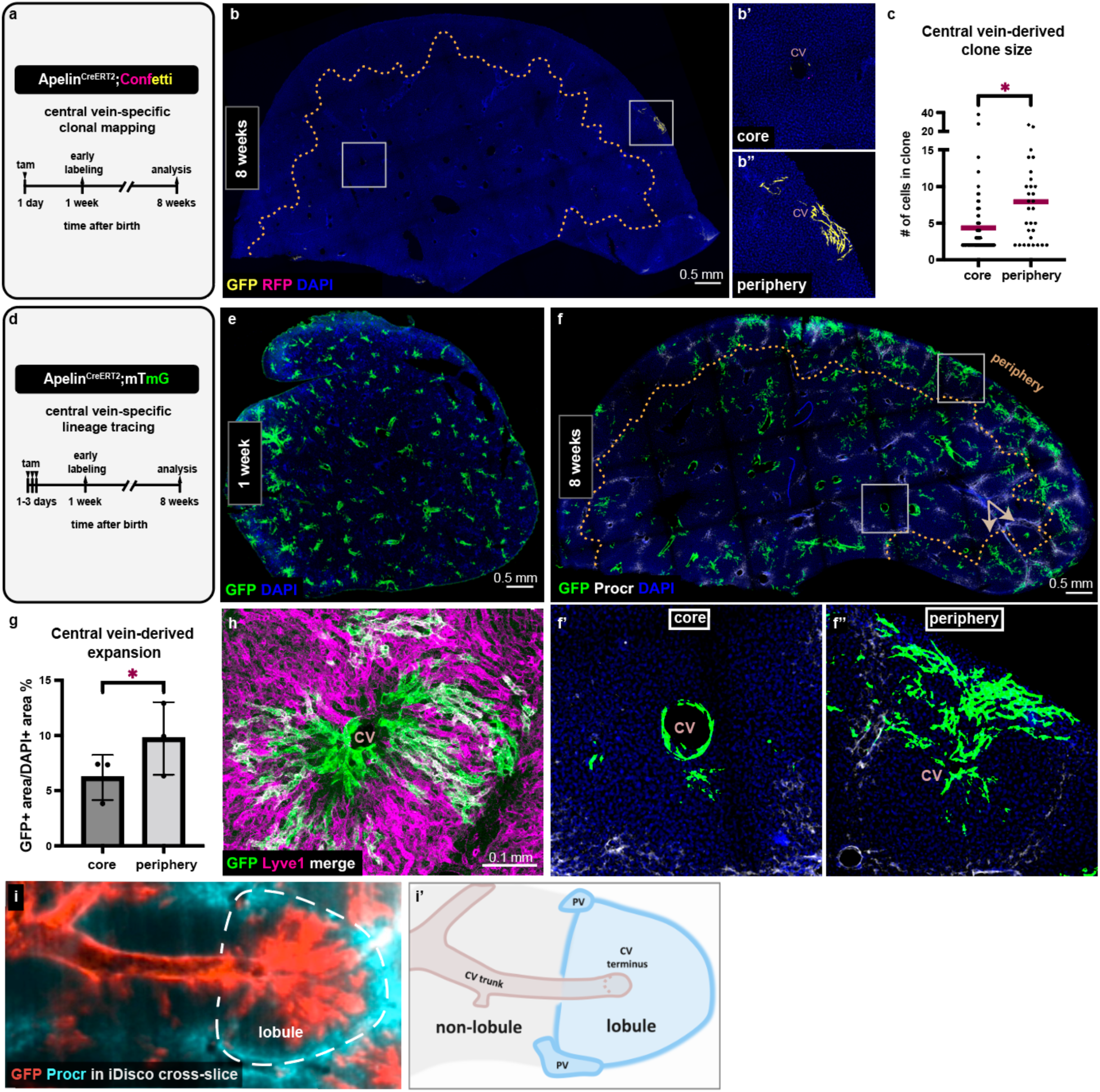
Central vein is the origin of new lobule vasculature at the periphery of the liver. **a**, Schematic summary of the experimental design to assess central vein-derived clonal expansion. **b-b’’,** Apelin^CreERT2^;Confetti thick liver section immunostained for GFP and RFP to mark the clones at 8 weeks. The dotted line delineates the periphery. **c,** Quantification of Apelin^CreERT2^;Confetti clone size measured as the number of cells per clone in the periphery compared to the core in 8-week-old mice. n=70 clones in the core and n=30 clones in the periphery from 3 mice were scored. Red bars represent the mean values. Welch’s t test, *p-value = 0.0108*. **d,** Schematic summary of the experimental design to assess central vein-specific lineage tracing. **e,** Apelin^CreERT2^;mTmG thick liver section immunostained for GFP to visualize the early labeling at 1 week. **f-f’’,** Apelin^CreERT2^;mTmG thick liver section immunostained for GFP and Procr to visualize the tracing at 8 weeks. The dotted line delineates the periphery. Arrows show two neighboring lobules with dramatically different amounts of expansion. **g,** Quantification of central vein-derived expansion in 8-week-old Apelin^CreERT2^;mTmG livers in the periphery compared to the core. n=3 mice, mean values ± s.d. Ratio paired t test, *p-value = 0.02 90*. **h,** Representative Apelin^CreERT2^;mTmG thick liver section immunostained for GFP and the sinusoidal marker Lyve1 to visualize the outcome of tracing in sinusoids. **i,** Cross-slice of iDisco-cleared Apelin^CreERT2^;mTmG whole lobe immunostained for GFP and Procr. The dotted line delineates lobule boundaries. **i’,** Cartoon representation of lobule and non-lobule arrangements. Central vein lineage tracing outcome is not represented for clarity. DAPI in blue depicts nuclei. tam, tamoxifen. CV, central vein. PV, portal vessel.

To test whether central veins indeed vascularize new lobules, we labeled all central veins neonatally using the mTmG reporter (Apelin^mTmG^) and traced the vein compartment as a whole until adulthood (Fig. 4d-f). Remarkably, endothelial cells sprouted and expanded out from the vein into the surrounding lobule in most lobules (79 %) (Fig. 4e-f, Extended Data Fig. 5c). Central vein cell expansion was most pronounced in peripheral lobules; here, the progeny frequently reached the lobule boundaries (Fig. 4f’-g). Since the lobule vascular bed is mainly composed of sinusoids^13,23^, this implied that the sprouting vein cells differentiate into sinusoids. To determine whether that is the case, we co-labeled Apelin^mTmG^ livers with the hepatic sinusoid marker Lyve1^23^. Indeed, the vein cell progeny formed the sinusoids of the lobule (Fig. 4h). These results identify the central vein as the vascular origin of newly forming lobules at the periphery.

Interestingly, each new lobule was vascularized as an independent unit. Every Apelin^Confetti^ clone examined was entirely contained within the same lobule as the vein of origin and did not cross over to neighboring lobules (Fig. 4b, b’’, Extended Data Fig. 5d). Furthermore, the extent of expansion was independent in each lobule. In two neighboring lobules, for instance, the vein of one lobule exhibited major expansion while the vein of the other expanded minimally (Fig. 4f, arrows). These data indicate that central vein cells sense lobule boundaries as they expand and solely vascularize their respective lobule. These findings suggest that central vein cell expansion can serve as a proxy for growth in a lobule.

If vein cell expansion is a suitable proxy for lobule growth, the expansion should be restricted to lobules and not occur in regions of the organ without proper lobule arrangement. To test this idea, we performed iDisco and whole lobe imaging of Apelin^mTmG^ livers (Extended Data Video 3). Volumetric imaging revealed vein cell expansion throughout the liver, distinguishing the regions with and without lobule arrangement. In lobules, the central vein terminus was seen crossing through the center, with cross-sections of portal vessels at the boundaries (Fig. 4i-i’). By contrast, in regions lacking lobule arrangement, central vein trunks were seen running longitudinally (Fig. 4i-i’). Strikingly, the expansion from central veins was restricted to lobules and did not occur in the non-lobule regions (Fig. 4i-i’). These data reinforce the suitability of central vein cell expansion as a proxy for lobule growth.

To rule out potentially rare but significant expansion from portal vessels that could contribute to new lobules, we also performed lineage-tracing of portal vessels using Efnb2^mTmG^. This was particularly important given the low frequency of clone induction in Efnb2^Confetti^ mice (see Methods). Efnb2^mTmG^ tracing studies showed minimal expansion of the cells out of their original portal vessel compartment and no biased expansion in peripheral lobules (Extended Data Fig. 5e-f). Thus, central veins, and not portal vessels, serve as the source of newly forming lobule vasculature. These results establish central vein cells as a source and a proxy for lobule vascularization and growth.

### Central vein cell sprouting declines rapidly during the neonatal period

The number of lobules that form in a given liver is expected to set the final size of the organ. Our early quantification showed an increase in the number of mature lobules after the first week (Fig. 1g). This suggested addition of new mature lobules but did not reveal when new lobules initiate or cease to initiate. To identify when new lobules cease to initiate and finalize their number, we monitored central vein cell sprouting as a proxy for new lobule initiation and asked at what age this process ceases.

We labeled central vein cells in one day-versus one week-old pups and compared their respective expansion by adulthood (Fig. 5a-f). This revealed a striking difference. When labeled at one week of age, central vein cells expanded minimally by adulthood, occupying only a small fraction of the periphery (Fig. 5c). This was in stark contrast with widespread expansion in peripheral lobules when central vein cells were labeled at one day of age (Fig. 5e-f). This difference may be caused either by major cellular expansion within the first week itself, or a decline in initial sprouting of the cells by one week of age. We assessed cellular expansion within the first week and found it to be minimal (Extended Data Fig. 4b). These results suggest a rapid decline in the initial sprouting of central vein cells within the first postnatal week.

**Fig. 5.**
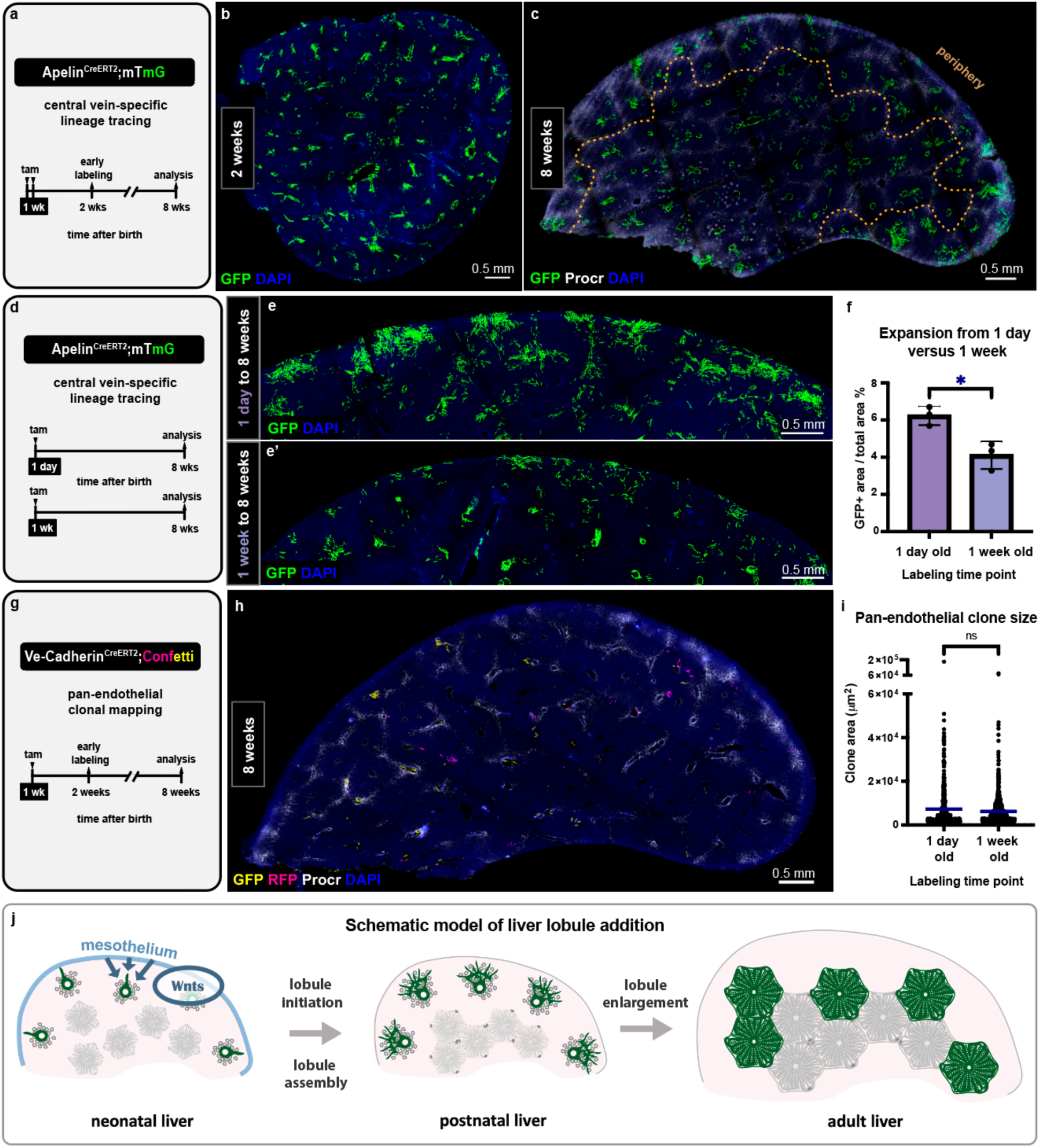
New lobule vascularization ceases to initiate neonatally. **a**, Schematic summary of the experimental design to assess central vein lineage tracing induced at 1 week (injections on postnatal days 6 and 7). **b,** Apelin^CreERT2^;mTmG thick liver section immunostained for GFP to visualize the early labeling at 2 weeks. **c,** Apelin^CreERT2^;mTmG thick liver section immunostained for GFP and Procr to visualize the tracing from 1 week to 8 weeks. The dotted line delineates the periphery. **d,** Schematic summary of the experimental design to compare central vein lineage tracing induced at 1 day (injections at postnatal days 1-3) versus 1 week (injections on postnatal days 6 and 7). **e-e’,** The periphery of Apelin^CreERT2^;mTmG livers lineage traced from 1 day-to-8 weeks in **e** and 1 week-to-8 weeks in **e’** immunostained for GFP. **f,** Quantification of central vein-derived expansion from 1 day-to-8 weeks compared to 1 week-to-8 weeks. n=3 mice per group, mean values ± s.d. Unpaired t test, *p-value = 0.0146*. **g,** Schematic summary of the experimental design to map endothelial cell expansion from 1 week on. **h,** VE-Cadherin^CreERT2^;Confetti thick liver section immunostained for GFP, RFP, and Procr to spatially map the clones that formed from 1 week to 8 weeks. **i,** Comparison of endothelial clone size in 8-week-old mice that were traced from 1 day versus from 1 week. n=457 clones from 3 mice (labeled at 1 day) and n=702 clones from 3 mice (labeled at 1 week). Blue bars represent mean values. Welch’s t test, *p-value = 0.0909*. **j,** Schematic model of the multicellular scale mechanisms that determine liver size. DAPI in blue depicts nuclei. ns, not significant; tam, tamoxifen.

We next sought to determine whether the drastic temporal change in sprouting is specific to vein endothelial cells, as opposed to occurring across endothelia. We performed Vecad^Confetti^ labeling in one week old pups and asked how the clonal expansion by adulthood compares to expansion from one day of age in bulk endothelial cells. We found that the average clone size as well as the clone size distribution are highly similar between those monitored from 1 day versus 1 week of age (Fig. 5g-i, Extended Data Fig. 6a-b). Thus, the majority of endothelial cells do not exhibit a drastic change in expansion behavior over the first postnatal week, and the rapid decline in sprouting is unique to central vein cells.

These findings suggest that the progenitor behavior of central vein cells is restricted to the neonatal period. The progeny of central vein cells continues to expand more at the periphery after the neonatal period (Extended Data Fig. 6c). This ensures vascularization of the growing peripheral lobules through the extended postnatal period.

Together, these results identify an endpoint for new lobule initiation within the neonatal week. Notably, this timing precedes the decline, or even the peak, of cell proliferation in the postnatal liver^24^ (Fig. 2a). Thus, vein cell sprouting emerges as a limiting factor in new lobule addition to the organ and in determining the adult liver size.

## Discussion

Our results identify five steps of multicellular integration that determine the mouse liver size. At birth, the liver is immature with no proper units or lobules (Fig. 5j). Over the first week after birth, the liver tissue is assembled into lobules (*lobule assembly*). Meanwhile, new lobules initiate at the organ edge, vascularized by sprouting of endothelial progenitors at central vein endings (*new lobule initiation*). Newly forming lobules obey pre-existing lobule boundaries. By the second week, vascular progenitors stop sprouting and new lobules cease to emerge, finalizing the lobule number (*conclusion of lobule initiation*). Lobules at the edge continue to grow through proliferation of hepatocytes and endothelial cells (*lobule enlargement)*, until all lobules reach proper size by weaning age (*conclusion of lobule enlargement*). At weaning, lobule number and size have been finalized, and the organ reaches its stereotyped size in ratio to the body (Fig. 5j).

These findings extend previous work that has established the neonatal period as the endpoint of functional unit addition in other rodent organs^25,26^. We therefore conclude that the neonatal time is a decisive period for rodent organ growth and size control. We find that the single cell-layered mesothelium encasing the liver spatially organizes growth during this period and ensures that the liver will reach its stereotyped size. The mesothelium encapsulates most vertebrate internal organs^27^, warranting further investigation into the role of this cell layer in organ growth and size control. At the center of this work, our results identify the hepatic vein as the *in vivo* vascular origin of new functional units in the mouse liver. Together with previous work^28–30^, our findings provide a foundation for building vascularized liver functional units with *bona fide* organization and size for regenerative medicine applications.

Finally, we demonstrate that the vascular source of new functional units is lost prior to the decline in cell proliferation in the liver. This implies that liver growth is limited by expansion of the vascular network rather than hepatic cell proliferation. Theoretical studies have previously proposed vascular geometry as a key determinant of allometric scaling in biology^3^. According to this model, the scaling of body parts with organism size can be explained by the features of branching transport systems, the vascular system in mammals. Our experimental results advance this theoretical work and lead us to postulate that temporally limited vascular expansion in mice helps scale liver size with organism size. Overall, this study provides a unique framework that we expect will guide future investigations into organ growth and size control.

## Materials and Methods

### Mice

The Institutional Animal Care and Use Committee at Stanford University approved all animal methods and experiments. Animal experiments were performed with mouse pups, and adult male and female 8-week-old mice. Data from males and females were pooled unless otherwise noted below or in the figure legends. Weaning age was set at 22-25 days. All mice were housed under a 12 h:12 h light:dark cycle at 22–25 °C and 30–70% humidity. Mice received rodent chow and water ad libitum. C56BL/6 mice were used for wild type studies. *mTmG* (007676), *Confetti* (017492), and *WT1^CreERT^*^2^ (010912) mice were purchased from The Jackson Laboratory and bred in house. *Tbx3^CreERT^*^2^ was generated by our lab^31^. *VE-Cadherin^CreERT^*^2^, also called *Cdh5-PAC-CreERT2 or* Cdh5-cre/ERT2^1Rha^, mice were generated by the Adams Lab^32^. *Wntless^flox^* mice were a gift from Alan Cheng and were generated by the Lang Lab^33^. *Apelin^CreERT^*^2^ mice were generated by the Zhou lab^34^ and have been used as an invaluable tool to target sprouting endothelial cells^35–39^, in line with identification of Apelin as an angiogenic factor and marker^40–42^. *Lyve1^CreERT^*^2^ mice were a gift from Darci Fink and were generated by the Tempero Lab^43^. *Esm1^CreERT^*^2^ mice were a gift and were generated by the Adams Lab^44^. *APJ^CreERT^*^2^ mice were generated by the Red-Horse lab^45^.

*Efnb2^CreERT^*^2^ mice were generated as part of this study, by Kyle Loh, and the Stanford Transgenic, Knockout, and Tumor Model Center. In these mice, the endogenous *Efnb2* gene was edited to replace the *Efnb2* stop codon with a *GSG-P2A-CreERT2-F5* cassette. A GSG-P2A linker was chosen, owing to the high translational skipping efficiency afforded by this linker. A single F5 site was also inserted downstream of *CreERT2*. This approach preserves the coding sequence of the endogenous *Efnb2* gene. These mutant mice were maintained as heterozygotes on a C57BL/6 background.

CreERT2 mice were crossed into mTmG or Confetti mice for lineage tracing and clonal mapping studies. To study Wntless loss of function in the mesothelium, WT1^CreERT2^ mice were first crossed into Wntless^flox/flox^ mice. Then, WT1^CreERT2^; Wntless^flox/+^ or WT1^CreERT2;^ Wntless^flox/flox^ mice were crossed into Wntless^flox/flox^ mice. Wntless^flox/+^ and Wntless^flox/flox^ littermates were used as controls. For *Apelin^CreERT^*^2^ lineage tracing studies, only males were used in order to avoid mosaic X chromosome inactivation, as the *Apelin* gene is X-linked.

### Tamoxifen treatment and lineage-tracing studies

Pups from Confetti or mTmG breedings were induced with Tamoxifen by intragastric injection before postnatal day 3 and by intraperitoneal injection afterwards. The doses and timing of injection were as follows: *Tbx3^CreERT^*^2^ *x Confetti* at postnatal day P1 (0.1 mg/ml)*, VE-Cadherin^CreERT^*^2^ *x Confetti* at P1 (0.067 mg/ml), *VE-Cadherin^CreERT^*^2^ *x Confetti* at P7 (0.1 mg/ml), *Apelin^CreERT^*^2^ *x Confetti* at P1 (2 mg/ml), *Efnb2^CreERT^*^2^ *x Confetti* at P1 (2 mg/ml), *Apelin^CreERT^*^2^ *x mTmG* at P1, P2, P3 (2 mg/ml each), *Apelin^CreERT^*^2^ *x mTmG* at P6, P7 (2 mg/ml each), *Efnb2^CreERT^*^2^ *x mTmG* at P1, P2, P3 (2 mg/ml each), *Lyve1^CreERT^*^2^ *x mTmG* at P1, P2, P3 (2 mg/ml each), *APJ^CreERT^*^2^ *x mTmG* at P1, P2, P3 (2 mg/ml each), *Esm1^CreERT^*^2^ *x mTmG* at P7 (2 mg/ml each). Tamoxifen was dissolved in sterile corn oil with 10 % ethanol. One week following Tamoxifen induction, tissues were collected for initial labeling analysis based on previous studies^46^.

### Vibrating microtome sectioning

Thick section processing and immunostaining protocol was adapted from Snippert *et al*, 2011^47^. For embedding and sectioning, Precisionary compresstome equipment and tools were used (VF-310-0Z). Briefly, livers were isolated and fixed in 4 % paraformaldehyde overnight at 4°C followed by PBS washes (3 × 10 min). For sectioning adult samples and postnatal samples older than one week, the whole left median lobe or half of the left liver lobe was embedded curvature down. The embedding was performed in the manufacturer’s mold in 4 % ultrapure low melting agarose prepared in PBS. For the sectioning of livers from younger pups, the whole left median lobe was embedded floating over a microcentrifuge tube cap in the mold to obtain proper agarose levels for reliable sectioning. The samples were left in the embedding mold for 5 minutes followed by incubation in the provided ice block for 10 minutes. The adult samples and postnatal samples older than one week of age were sectioned at 100 µm, while samples from younger pups were sectioned at 50 µm. All sectioning was carried out at speed set to 3 and vibration set to 4.

For immunostaining, the compresstome sections were permeabilized in PBS containing Triton X-100 0.1% for 1 hour, blocked in PBS with Triton X-100 0.1% containing normal donkey serum at 5 % for 1 hour, and incubated in primary antibodies listed in the antibodies section for 2 overnights at 4°C. The sections were then washed in PBS containing Triton X-100 0.1% (4 x 15 min) and incubated in secondary antibodies raised in donkey conjugated to Alexa Fluor 488, 555, or 647 (1:400; Jackson Immunoresearch Labs) overnight at 4°C. Next, the sections were washed in PBS containing Triton X-100 0.1% (4 x 15 min), incubated in DAPI solution in PBS (1:10000, cat no) for 30 minutes, and washed again for 15 minutes. Lastly, using a paintbrush, the sections were mounted on a slide with Prolong Gold (with DAPI) and protected with a coverslip.

### Confocal imaging

Immunostained, thick fluorescent liver sections were imaged using a Leica SP8 white light laser confocal. All images were acquired at 10X magnification, except for those used for the hepatocyte clonal analysis which were acquired at 20X magnification. Images of full liver sections were tiled and stitched by automatic merge using the Navigator view on the LasX Software (3.5.5.19976). Z stacks of images were projected into a 2-dimensional image by applying the maximum intensity projection on FIJI (ImageJ).

### Volumetric Immunofluorescence

Whole left median lobes were immunostained using a protocol adapted from the previously published iDisco protocol without Methanol pre-treatment^14^. The livers were perfused with fixative at the time of collection for thorough fixation. The perfusion was carried out by inserting a catheter connected to a perfusion pump through the inferior vena cava. Each liver was perfused with 10 ml PBS followed by 6 ml 4 % paraformaldehyde. The samples were then fixed further in 4 % paraformaldehyde overnight at 4°C and washed in PBS (3 × 10 min). For iDisco pre-treatment, the samples were washed in PBS containing 0.2% TritonX-100, referred to as PTx.2 (1 hour x 2), incubated in PTx.2 with 20 % dimethyl sulfoxide overnight at 37°C and then in PBS with 20 % dimethyl sulfoxide containing a combination of detergents (0.1% Tween, 0.1% TritonX-100, 0.1% Deoxycholate, 0.1% NP40) overnight at 37°C. The samples were washed in PTx.2 (1 hour x 2) to complete the pre-treatment. For immunolabeling, the samples were incubated in permeabilization solution (PTx.2 containing 20 % dimethyl sulfoxide and 2.3 % glycine) for 2 days at 37°C, in blocking solution (PTx.2 containing 10 % dimethyl sulfoxide and 6 % normal donkey serum) for 2 days at 37°C, and in primary antibodies for a week at 37°C. The primary antibodies were prepared in PBS containing 0.2 % Tween20 and 10 µg/ml, referred to as PtwH, with 5 % dimethyl sulfoxide and 3 % normal donkey serum. The samples were then washed in PtwH (5 x 1 hour) and left washing overnight. Next, the samples were incubated in secondary antibodies (prepared in PtwH containing 3 % normal donkey serum) for a week at 37°C followed by washes (5 x 1 hour) and an overnight incubation in PtwH. The secondary antibodies used were raised in donkey and conjugated to Alexa Fluor 488, 555, or 647 (1:400; Jackson Immunoresearch Labs). These antibodies were centrifuged prior to diluting to avoid aggregates. For clearing, the samples were dehydrated in a methanol series with increasing concentration for 1 hour each (20 %, 40 %, 60%, 80%, 100% x 2). The samples were left in 100 % methanol overnight, and then incubated in 2:1 dichloromethane: methanol solution for 3 hours. Finally, the samples were incubated in 100 % dichloromethane for 15 minutes twice to rinse the methanol and transferred into dibenzyl ether in a completely filled tube. Except for this last step, all incubations were carried out in 4.5 ml solution volume. 5 ml Eppendorf tubes were used. For livers from 1 week old pups, the samples were embedded in 1 % agarose prepared in TAE buffer prior to clearing.

### Light sheet imaging

The whole liver lobe samples were imaged on a Miltenyi / LaVision Biotec Ultramicroscope II. Volumetric images were generated using the Imaris (Bitplane) software, version 10.0.0. Deconvolution was performed for improved visualization of volumetric images.

### Antibodies

The following primary antibodies were used at indicated concentrations for the immunostaining of compresstome sections or iDisco whole lobe samples: goat anti-EPCR (R&D Systems, AF2749, 1:100), rat anti-Endomucin (Santa Cruz Biotech, sc-65495, 1:100), rabbit anti-phospho histone H3 (Milipore Sigma, 06-570, 1:100), chicken anti-GFP (Abcam, ab13970, 1:500), rabbit anti-RFP (Rockland, 600-401-379, 1:200), and rabbit anti-Lyve1 (Novus Biologicals, NB600-1008, 1:500).

### In situ hybridization

In situ hybridization on postnatal liver samples was performed using the high-sensitivity RNA amplification and detection technology RNAscope (Advanced Cell Diagnostics). The samples were treated according to the manufacturer’s instructions. Briefly, the samples were fixed in 10 % neutral buffered formalin overnight at room temperature. The samples were paraffin-embedded and sectioned within three days prior to and baked the day before in situ hybridization. The RNAscope chromogenic 2.5 HD Duplex Detection Kit was used. The antigen retrieval step was performed for 12 minutes. The signal detection steps were performed for 30 minutes. The used probes are as follows: Mm-Procr-Ch2 (410321-C2), Mm-Tbx3-Ch1 (438211), Mm-WT1-Ch2 (432711-C2), and Mm-Wntless-Ch1 (405011).

### Image processing, analysis, and quantification

Tiled and stitched full-section confocal images of z-stacks (30-50 um) were converted into maximum projection images, and the projected images were used for further analyses unless otherwise noted below. Whole left median lobes were used all experiments unless otherwise noted below or in the figure legends. Same thickness images were used across samples for quantification of cell numbers or the comparison of fluorescence intensity. Images used for comparison were acquired with the same imaging settings. For DAPI normalization, DAPI-stained area was measured following intensity thresholding. For the presentation of Procr and GFP immunostaining in figures, the images were despeckled in FIJI (ImageJ). On full liver section images in Fig. 1c, 2i, 2l, and Extended Data Fig. 4d, autofluorescence signal due to trapped air bubbles outside of the tissue section was masked during figure assembly for presentation purposes. Image quantifications are described in detail below. Unless individual values were plotted as denoted in the figure legend, the average value derived from 2-3 sections of each sample was calculated and presented as a data point.

#### Lobule size quantification

Procr-immunostained full liver section images were cropped into single lobule size for all lobules identified. 140-210 lobules from a total of 3 sections per sample and from 3 mice at each age were analyzed. In Wntless^meso^ litters, 28-32 lobules from a total of 3 sections per sample and from 2 mice were analyzed for each genotype. The images were run through a custom MatLab script generated in this study to assess the number of nuclei per lobule.

#### Quantification of lobule numbers

Volumetric iDisco samples immunostained for Procr were used for the quantification of mature lobules in the left median lobe. The lobules were manually counted on FIJI (ImageJ). The mature lobule was defined as one with boundaries marked by Procr immunostaining and a central vein lumen which can be identified against the background tissue signal. It was empirically determined that lobule numbers quantified on sections accurately represents the total number of lobules when two samples are being compared, and thus the comparison of lobule numbers in Wntless^meso^ litters was performed on sections. 2-3 sections from 4 Wntless^meso^ and 2 control littermates were analyzed.

#### Edge measurements at neonatal ages

On Procr-immunostained full liver section images, the Procr-positive portal vessel structures closest to the edge of the section were manually traced and connected. This tracing partitioned a thin layer of tissue that constitutes the edge and the rest of the tissue that constitutes the core.

#### Periphery measurements at adulthood

On Procr-immunostained full liver section images, the lobule boundaries were manually traced at one lobule distance along the outer edge of the liver lobe, counting three lobules up from the corner of the distal tip. In cases where the lobule hexagon is not complete at the section edge, it was counted as a lobule when the corresponding central vein is discernable.

#### Quantification of mitosis

Phospho-histone H3 immunostained full liver section images were used. In FIJI (ImageJ), intensity thresholding and size filtering were applied to filter out noise. The same thresholds were used across samples under comparison. The “analyze particles” function of FIJI (ImageJ) was applied to the post-filtering images to identify the nuclei with pHH3 staining. The obtained numbers were normalized to the tissue area defined as the DAPI stained area. The normalized numbers represent the mitotic or pHH3 index. Relative numbers were generated for plots by multiplying the DAPI-normalized ratio by 10^5^ or 10^6^ as denoted. To quantify the distribution of mitosis, a similar approach was used accompanied by periphery identification. The corresponding Procr-stained split image was used to identify the periphery as described above. The corresponding particle analysis image and the DAPI split image were both partitioned into the periphery and the complementary core in sync with the corresponding Procr image. The number of particles was normalized to the corresponding DAPI-stained area for the periphery and the core separately, and the DAPI-normalized ratios were multiplied by 10^5^ or 10^6^ as denoted to generate relative numbers for plots.

#### Hepatocyte clone size quantification

Hepatocyte clone size was assessed by manual cell counting on slices of z-stacks of RFP- and GFP-immunostained Tbx3^CreERT2^; Confetti samples in FIJI (ImageJ). 126 clones in the core and 41 clones in the periphery from a total of 3 sections collected from 3 mice were analyzed. Cells of the same color (RFP or GFP) and with the same reporter localization (nuclear, cytoplasmic, or membrane) within the empirically determined one-cell-distance were considered belonging to the same clone. Single cells were excluded from the analysis.

#### Endothelial clone size quantification

Endothelial clone size was assessed on maximum projection images of RFP-immunostained VE-Cadherin^CreERT2^; Confetti samples. 457 (labeled at 1 day old) and 702 (labeled at 1 week old) clones from a total of 2-3 sections per sample and from 3 mice at each age were analyzed. Single cells were excluded from the analysis. The area of each clone was measured using a MatLab script for endothelial clone area quantification generated in this study. Rarely, clones were erroneously called by the script due to high background in images and discarded post-data collection (0.6 % error rate). The clones were then binned by area measurement, and the number of clones in each bin were plotted to represent clone size distribution. The irregular right tail of the histogram was used to identify rare, large clones which were then defined as giant clones. For comparison of clone size between the core and the periphery, the same script was used in combination with Procr-guided periphery identification. Out of 457 clones, 128 localized to the periphery while 329 localized to the core. The periphery was manually traced using the corresponding Procr-stained split images as part of the MatLab script.

#### Labeling efficiency quantification

The labeling efficiency was measured by manually counting the number of labeling events on maximum projection images of RFP- and GFP-immunostained VE-Cadherin^CreERT2^; Confetti samples in FIJI (ImageJ). The number of labeling events was defined as the sum of the number of single cells and the number of clones where each clone is counted as one labeling event. This number was normalized to the tissue area defined as the DAPI stained area. Relative numbers were generated for plots by multiplying the DAPI-normalized ratio by 10^6^.

#### Quantification of central vein-derived clone size

The size of central-vein derived clones was assessed on maximum projection images of RFP- and GFP-immunostained Apelin^CreERT2^; Confetti samples. 70 clones in the core and 30 clones in the periphery from a total of 3 sections per sample and from 3 mice were analyzed. The cells were manually counted in FIJI (ImageJ). Single cells were excluded from the analysis (158 single cells out of 258 total labeling events).

#### Quantification of central vein-derived expansion

The extent of central vein-derived expansion was measured using intensity thresholding on maximum projection images of GFP-immunostained Apelin^CreERT2^; mTmG samples in FIJI (ImageJ). 1-3 sections from 3 mice at each age were analyzed. The GFP-stained area was normalized to the tissue area defined as the DAPI-stained area and plotted as a percentage. The number of veins with different central-vein derived expansion outcomes was manually evaluated on maximum projection images of GFP-immunostained Apelin^CreERT2^; mTmG samples in FIJI (ImageJ).

#### Quantification of portal vessel-derived clone size

The size of portal-vessel derived clones was assessed on maximum projection images of RFP- and GFP-immunostained Efnb2^CreERT2^; Confetti samples. Despite maximum Tamoxifen dosage, the labeling events in these mice remained very low. Assessment of 3 sections per sample from 3 mice yielded only 17 clones in the core and 8 clones in the periphery. The number of cells in these clones was manually assessed in FIJI (ImageJ). Single cells were excluded from the analysis (29 single cells out of 54 total labeling events).

#### Quantification of central vein labeling outcomes

The outcomes of central vein labeling were assessed manually on maximum projection images of RFP- and GFP-immunostained Apelin^CreERT2^; Confetti samples in FIJI (ImageJ). A total of 77 clones and 59 single cells from 1-3 sections per sample collected from 4 mice were analyzed.

#### Quantification of portal vessel-derived expansion

The extent of portal vessel-derived expansion was measured using intensity thresholding on maximum projection images of GFP-immunostained Efnb2^CreERT2^; mTmG samples in FIJI (ImageJ). 2 sections from 2 mice at each age were analyzed. The GFP-stained area was normalized to the tissue area defined as the DAPI-stained area and plotted as a percentage.

## Statistical analyses

Statistical analyses were performed using Prism 9 (Graphpad Software 9.5.1). Unpaired t test was applied to comparisons between two conditions with similar sample sizes except that of average sample readouts between the core and the periphery. The latter were analyzed using the ratio paired t test. When comparing two conditions with different sample sizes (difference > 50 %), Welch’s t test was performed to account for different variances between datasets. Ordinary one-way Anova analysis and Tukey’s multiple comparisons test was applied to comparisons across more than two conditions unless variances differed. Brown-Forsythe Anova analysis and Dunnett’s multiple comparisons test were applied to comparisons across more than two datasets with differing variances. All tests were two-sided. *P* < 0.05 was considered statistically significant. The significance annotations are * p-value < 0.05, ** p-value < 0.01, *** p-value < 0.001, **** p-value < 0.0001. Sample size was not predetermined with statistical methods. Previous studies using similar experimental approaches guided the decisions on sample size. The investigators were blinded during the treatment, collection, immunostaining, imaging, and quantification of Wntless^meso^ studies, and during the quantification of the number of lobules in wild types. The rest of the experiments were performed without blinding.

## Author contributions

D.B.A. and R.N. designed the experiments. D.B.A., K.P., S.R., T.A., and M.F. performed the experiments. D.B.A. and K.P. performed all experiments with the following exceptions: S.L.Z. and E.R. provided expertise in image analysis and generated Matlab scripts for the analyses of clone size and lobule size. S.R. performed the experiments outlined in Fig.2a-c. M.F. performed in situ hybridization experiments presented in Fig. 3a and Fig S3a-b. T.A. characterized the Tbx3^CreERT2^ mouse line and provided key conceptual input from the earliest steps of this work. K.L. generated the crucial Efnb2^CreERT2^ line for this work and provided key conceptual input throughout this study. K.R. provided conceptual input central to the progress of this work. D.B.A. and R.N. wrote the manuscript with input from all co-authors. All authors read and approved the manuscript for submission. R.N. supervised the study.

## Acknowledgements

We thank members of the Nusse lab, Arnaldo Carreira-Rosario, and John Vaughen for conceptual input for this work and critical review of the manuscript. We are grateful to Catriona Logan additionally for guidance and assistance with animal work and imaging. We thank Abby Sarkar for establishing and optimizing compresstome sectioning methods in the Nusse lab. We thank Gordon Wang for assistance with light sheet microscopy which was made available by the Stanford Wu Tsai Neuroscience Microscopy Core through the NIH shared equipment grant 1S10OD025091-01. We are grateful to Ralf Adams, Alan Cheng, Darci Fink, and Bin Zhou for generously sharing key mouse lines for this work.

---

**Extended Data Fig. 1.**
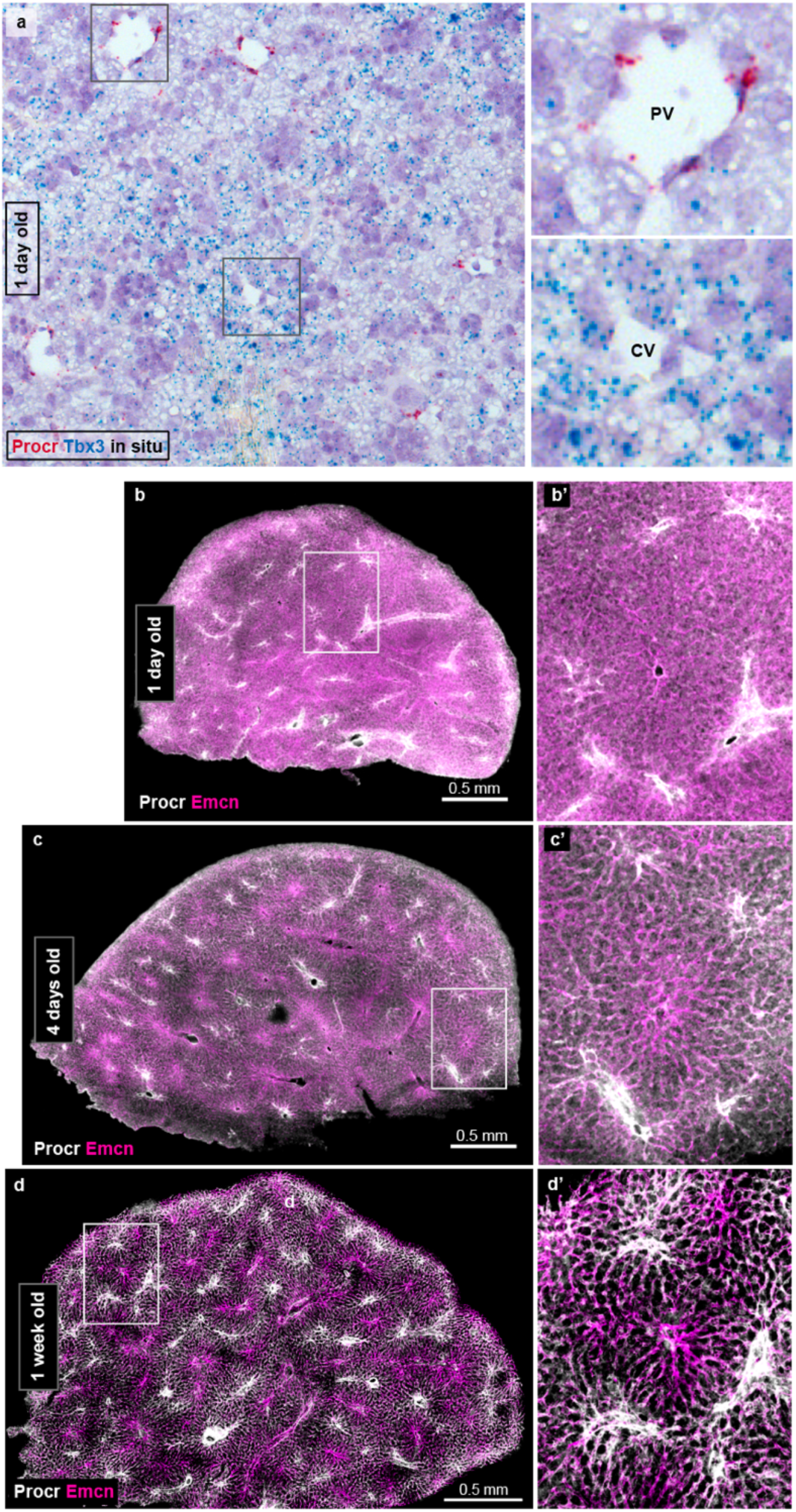
Temporal assessment of liver lobules. **a**, *In situ* hybridization on a representative wildtype liver section at 1 day old for the lobule boundary marker *Procr* and *T-box transcription factor 3 (Tbx3)* expressed in zone 2 and 3 hepatocytes. **b-d’,** Thick liver sections immunostained for the lobule boundary marker Procr and the venous marker Endomucin in 1-day-old, 4-day-old, and 1-week-old wildtype mice. PV, portal vessel. CV, central vein.

**Extended Data Fig. 2.**
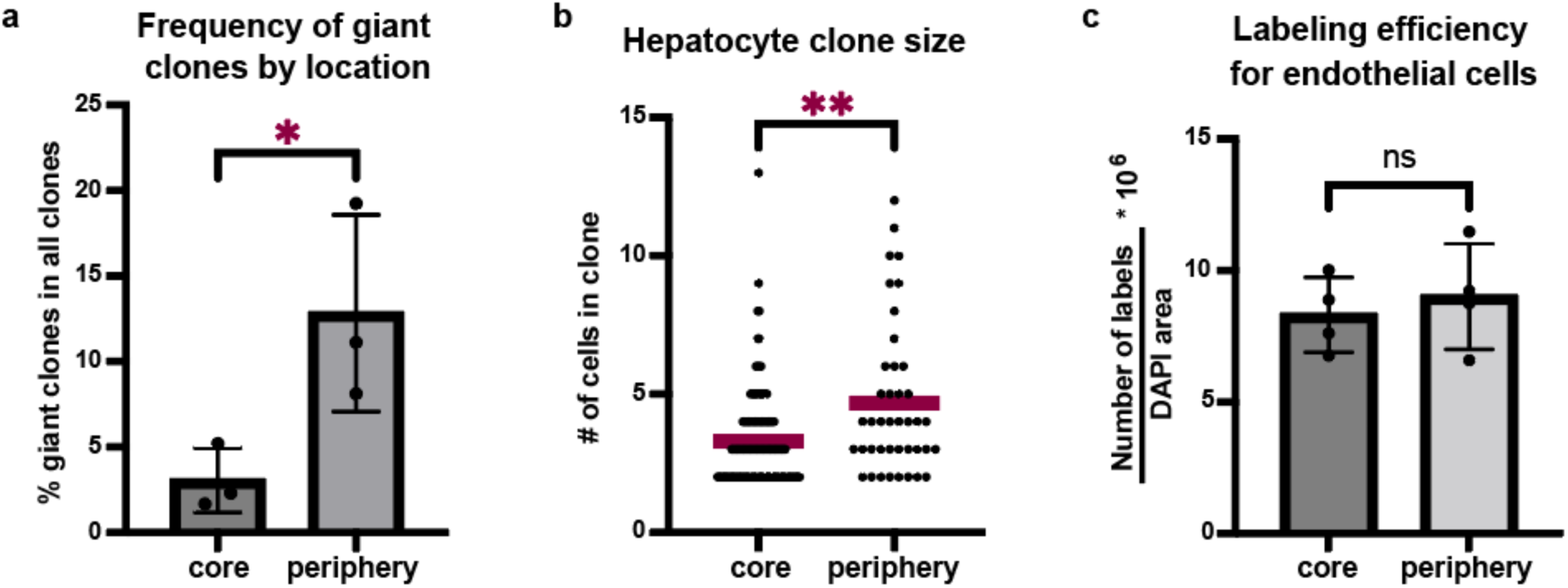
Clone size and labeling efficiency quantifications. **a**, Quantification of the frequency of giant clones (area > 30,000 μm^2^) in the periphery compared to the core. Unpaired t test, *p-value = 0.0493*. **b,** Quantification of hepatocyte clone size measured as the number of cells per clone in the periphery compared to the core in 8-week-old Tbx3^CreERT2^;Confetti livers. n=126 (core) and n=41 (periphery) clones from 3 mice. Red bars represent the mean values. Welch’s t test, *p-value = 0.0035*. **c,** Quantification of endothelial cell labeling efficiency measured as the number of labeling events normalized to the tissue area in 1-week-old VE-Cadherin^CreERT2^;Confetti livers. n=4 mice per group, mean values ± s.d. Unpaired t test, *p-value = 0.5933*. ns, not significant.

**Extended Data Fig. 3.**
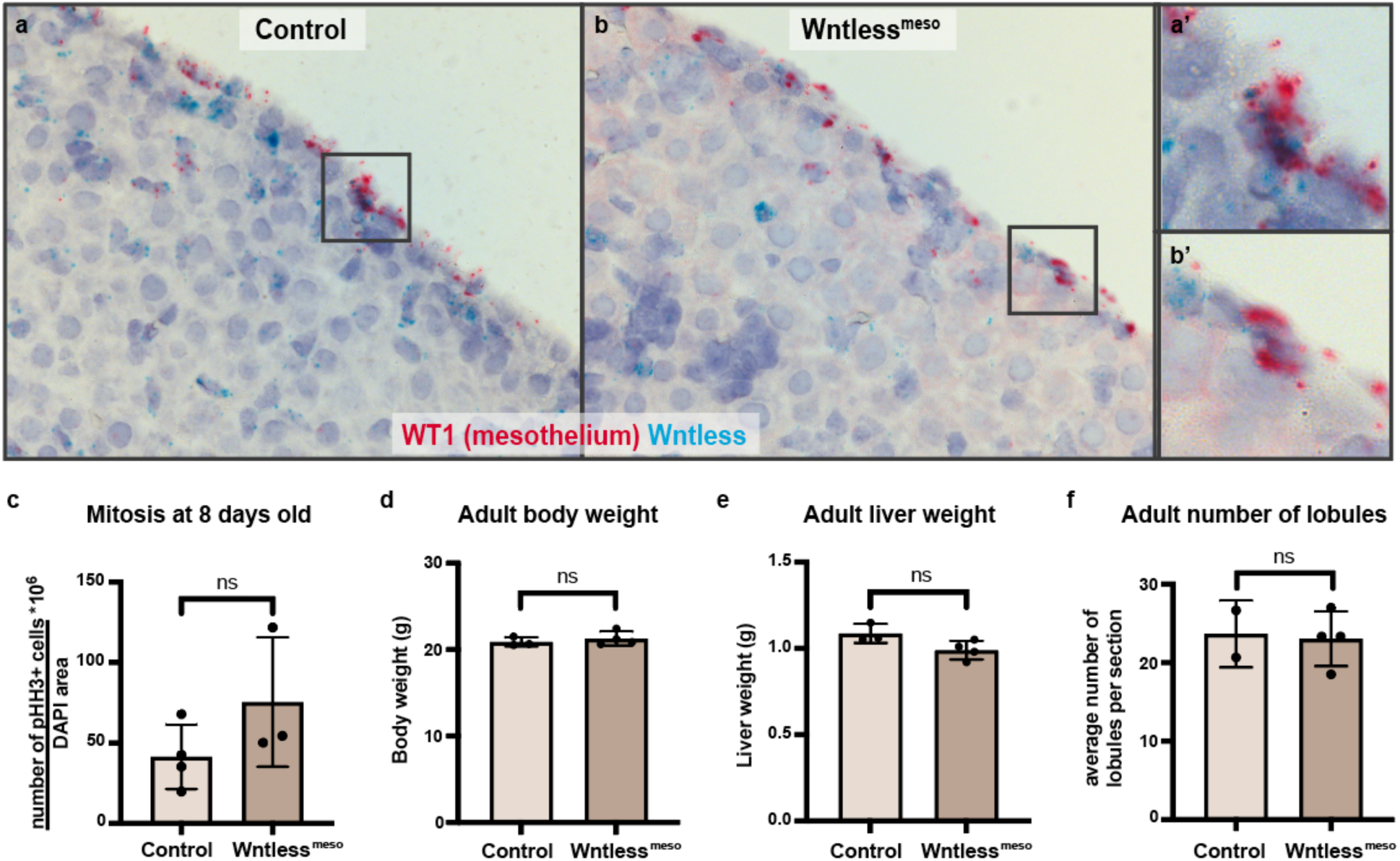
Characterization of mesothelial-specific Wntless loss of function in the liver. **a-b’,** In situ hybridization on liver sections for *Wntless* and the mesothelial-specific *Wilms Tumor 1 (WT1)* from 8-day-old control and mesothelial-specific Wntless knock-out (KO), or Wntless^meso^, mice. **c,** Quantification of overall mitotic index in 8-day-old control (n=4) and Wntless^meso^ (n=3) mice, mean values ± s.d. Unpaired t test, *p-value = 0.1954*. **d,** Body weight in 8-week-old control (n=3) and Wntless^meso^ (n=4) mice, mean values ± s.d. Unpaired t test, *p-value = 0.4955*. **e,** Absolute liver weight in 8-week-old control (n=3) and Wntless^meso^ (n=4) mice, mean values ± s.d. Unpaired t test, *p-value = 0.0693*. **f,** Quantification of the number of lobules in 8-week-old control (n=2) and Wntless^meso^ (n=4) mice, mean values ± s.d. Unpaired t test, *p-value = 0.8545*. ns, not significant.

**Extended Data Fig. 4.**
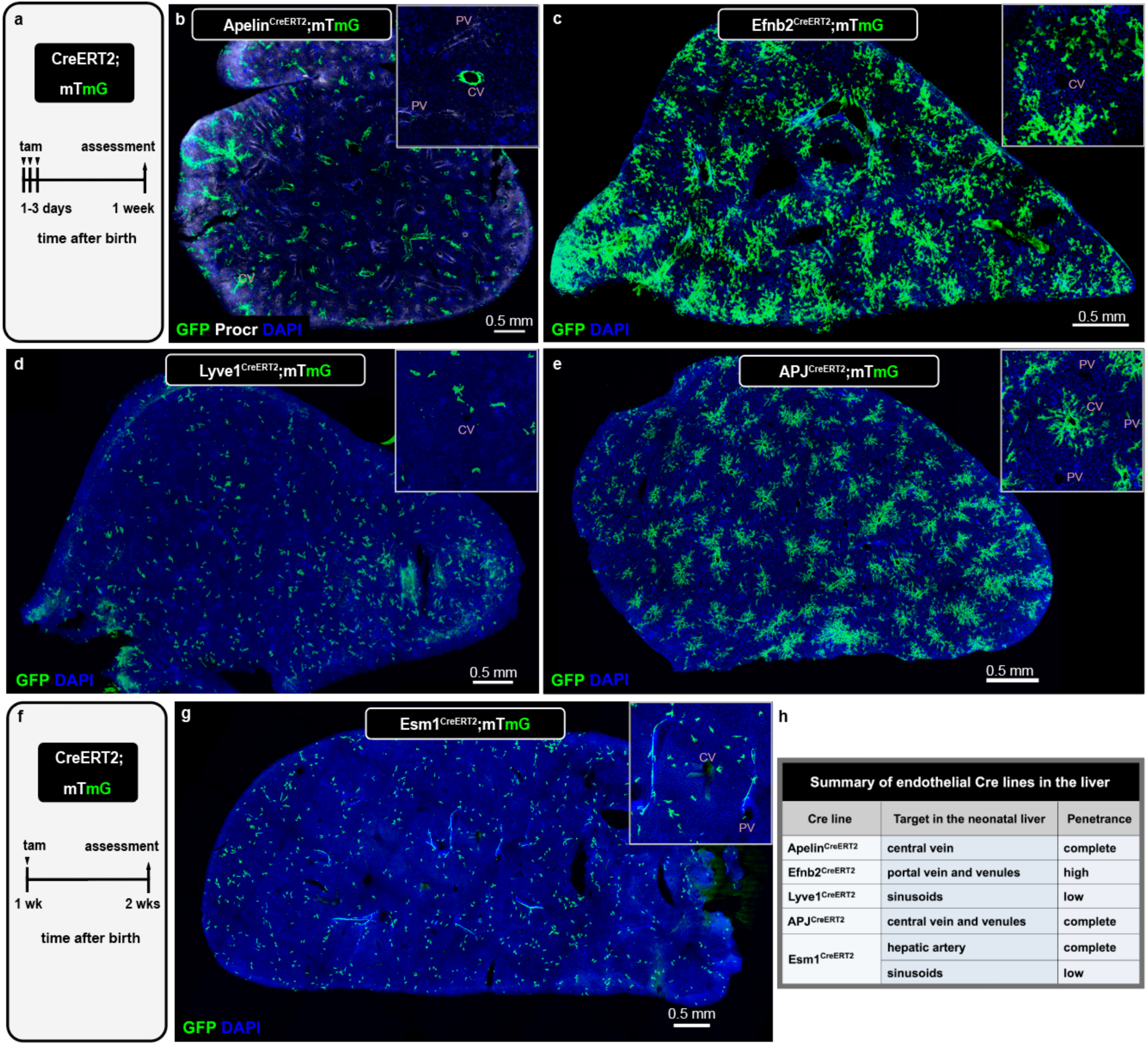
Genetic targeting of vessel subtypes in the liver. **a**, Schematic summary of the experimental design to label different vessel subtypes as a whole compartment using CreERT2 lines in the postnatal liver. **b,** Apelin^CreERT2^;mTmG thick liver section immunostained for GFP and the lobule boundary marker Procr at 1 week. **c-e,** Efnb2^CreERT2^;mTmG, Lyve1^CreERT2^;mTmG, and APJ^CreERT2^;mTmG thick liver sections immunostained for GFP at 1 week. **c,** Efnb2^CreERT2^;mTmG thick liver section immunostained for GFP at 1 week. **f,** Schematic summary of the experimental design to label vessels in Esm1^CreERT2^;mTmG mice. **g,** Esm1^CreERT2^;mTmG thick liver section immunostained for GFP at 2 weeks. h, Summary table of labeling outcomes for the vascular subtype Cre lines in the postnatal liver. DAPI in blue depicts nuclei. tam, tamoxifen. PV, portal vessel. CV, central vein.

**Extended Data Fig. 5.**
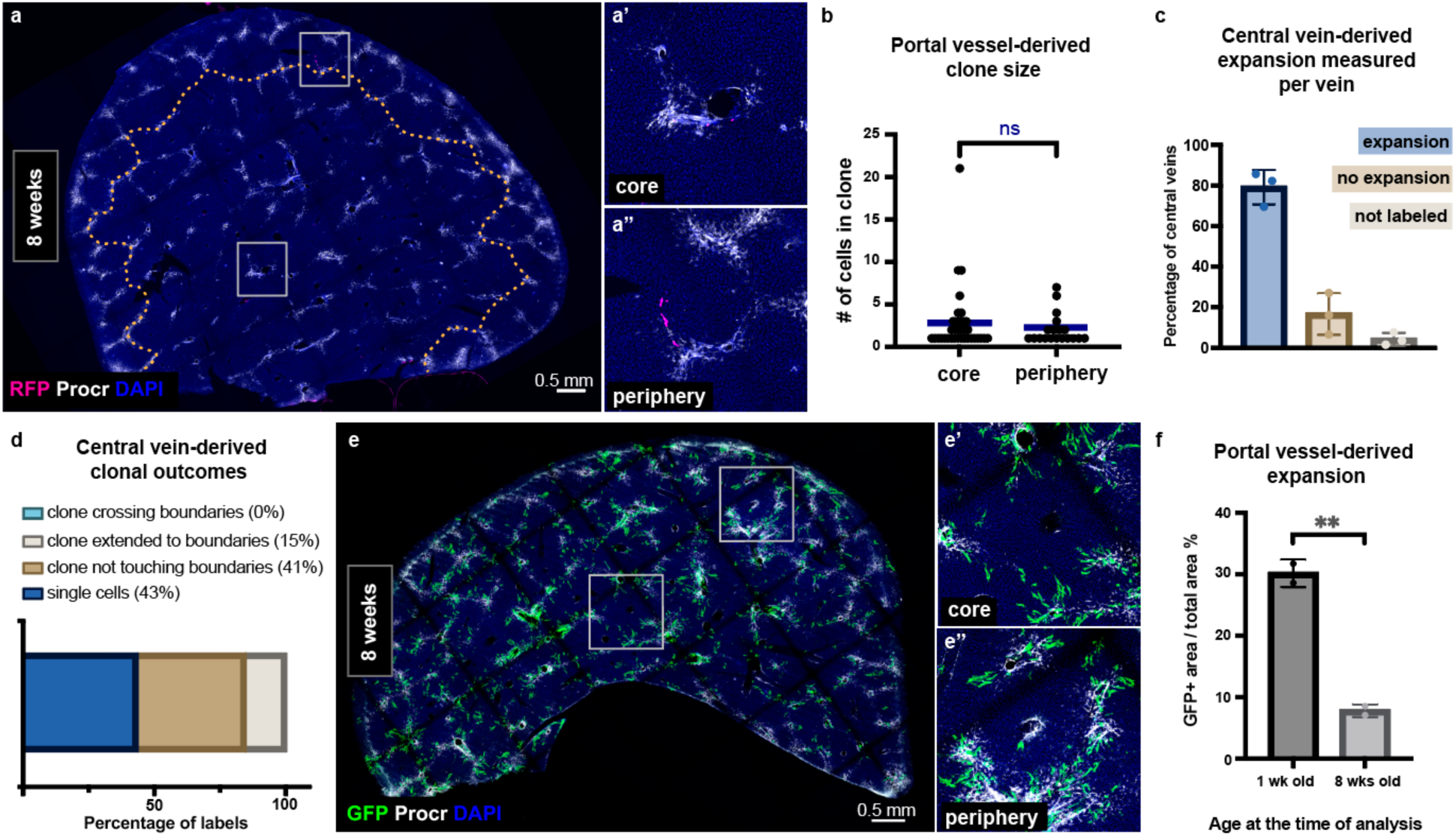
Contribution of portal vessels and the central vein to new lobule vasculature. **a-a’’,** Efnb2^CreERT2^;Confetti thick liver section immunostained for RFP to mark the clones and Procr to mark the lobule boundaries at 8 weeks following induction at postnatal day 1. The dotted line delineates the periphery. **b,** Quantification of portal vessel-derived clone size measured as the number of cells per clone in the periphery (n=8) compared to the core (n=17) from 3 Efnb2^CreERT2^;Confetti mice at 8 weeks. Blue bars represent mean values. Welch’s t test, *p-value = 0.3828*. **c,** Quantification of the percentage of central veins with different expansion or labeling behaviors (expansion at 79 %, no expansion at 17 %, and not labeled at 3 %) in 8-week-old Apelin^CreERT2^;mTmG mice. 637 central veins scored from n=3 mice. **d,** Summary graph of the percentage of central vein-derived clonal outcomes categorized by the spatial pattern of clones relative to lobule boundaries in 8-week-old Apelin^CreERT2^;Confetti mice. n=272 labels (clones and single cells) were scored from 4 mice. **e-e’’,** Efnb2^CreERT2^;mTmG thick liver section immunostained for GFP and the lobule boundary marker Procr at 8 weeks following induction at postnatal days 1-3. **f,** Quantification of portal vessel-derived expansion at 8 weeks compared to the early labeling at 1 week in Efnb2^CreERT2^;mTmG mice. n=2 mice per group, mean values ± s.d. Unpaired t test, *p-value = 0.0061*. ns, not significant. DAPI in blue depicts nuclei.

**Extended Data Fig. 6.**
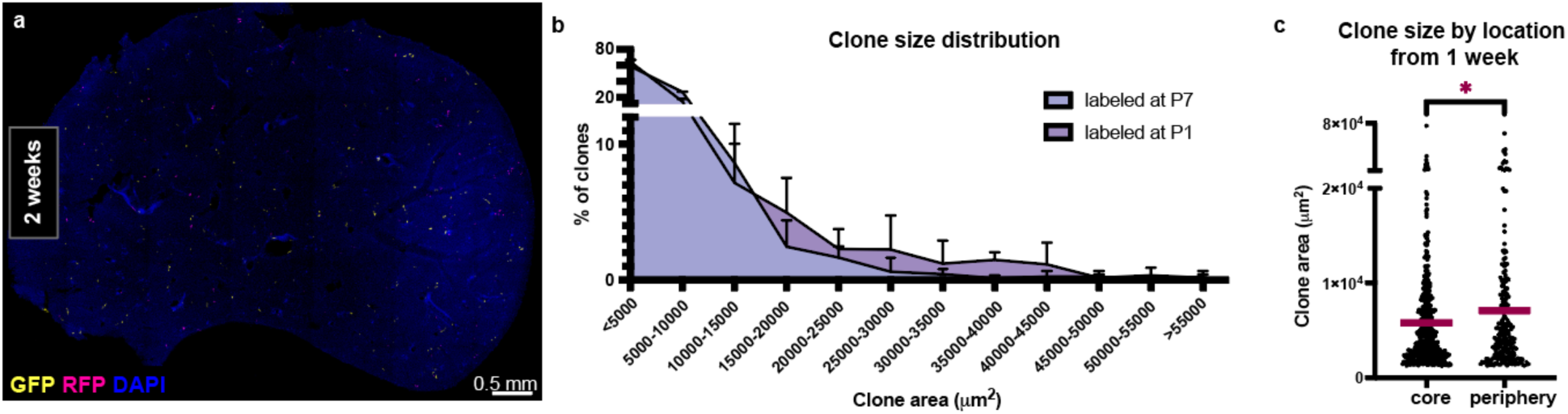
Endothelial expansion after the first neonatal week. **a**, VE-Cadherin^CreERT2^;Confetti thick liver section immunostained for GFP and RFP to mark the labeling events at 2 weeks following induction at 1 week. b, Distribution of endothelial clone size in 8-week-old VE-Cadherin^CreERT2^;Confetti livers comparing induction at 1 day versus 1 week, n=457 clones from 3 mice (induced at 1 day) and n=702 clones from 3 mice (induced at 1 week). Mean values + s.d. **c,** Quantification of clone size in the periphery compared to in the core from 8-week-old VE-Cadherin^CreERT2^;Confetti mice following induction at 1 week. n=473 (in the core) and n=229 (in the periphery) clones were scored from 3 mice. Red bars represent mean values. Welch’s t test, *p-value = 0.0455*. DAPI in blue depicts nuclei.

**Extended Data Video 1** Animated z-stack reconstruction of a whole lobe from an 8-week-old wild type mouse liver, imaged on a light sheet microscope following iDisco clearing and Procr immunostaining (white) to mark lobule boundaries. Lobules are seen covering the surface of the lobe. Lobule portal vessels and central veins connect to the larger vascular trees deeper into the lobe.

**Extended Data Video 2** Zoom in on a few lobules from the Extended Data Video 1, an animated z-stack reconstruction of an 8-week-old liver lobe that was iDisco-cleared, immunostained for Procr (white, lobule boundaries), and imaged on a light sheet microscope. A lobule with distinct boundaries is seen close to the surface of the lobe. Deeper into the organ, the boundaries of the lobule fade and the lobule arrangement disappears. The central vein of the lobule merges with the central vein of a nextdoor lobule, signifying the vein branching point. Many more branching points appear and lead to the trunk of the main hepatic vein deeper into the organ.

**Extended Data Video 3** Animated z-stack reconstruction of an 8-week-old Apelin^CreERT^^2^;mTmG whole liver lobe imaged on a light sheet microscope following iDisco clearing and immunostaining for Procr (cyan) to mark lobule boundaries and GFP (orange) to mark cellular expansion derived from central veins. Lobules are seen on the surface of the lobe with prominent vein cell expansion. Deeper into the organ, many central veins bend and start to run longitudinally, eventually feeding into larger vein branches. In these regions that lack lobule arrangement, vein cell expansion is virtually absent.

